# A Robust Model for Circadian Redox Oscillations

**DOI:** 10.1101/590570

**Authors:** Marta del Olmo, Achim Kramer, Hanspeter Herzel

## Abstract

The circadian clock is an endogenous oscillator that controls daily rhythms in metabolism, physiology and behavior. Although the timekeeping components differ among species, a common design principle is a transcription-translation negative feedback loop. However, it is becoming clear that other mechanisms can contribute to the generation of 24 h rhythms. In mammalian adrenal gland, heart and brown adipose tissue, peroxiredoxins (Prx) exhibit 24 h rhythms in their redox state. Such rhythms are generated as a result of an inactivating hyperoxidation reaction that is reduced by coordinated import of the sulfiredoxin (Srx) into the mitochondria. However, a quantitative description of the Prx/Srx oscillating system is still missing. We investigate the basic principles that generate mitochondrial Prx/Srx rhythms using computational modeling. We observe that the previously described delay in mitochondrial Srx import, in combination with an appropriate separation of fast and slow reactions is sufficient to generate robust self-sustained relaxation-like oscillations. We find that our conceptual model can be regarded as a series of three consecutive phases and two temporal switches, highlighting the importance of delayed negative feedback and switches in the generation of oscillations.

## 1. Introduction

The Earth’s regular 24 h rotation has led to the evolution of circadian oscillators in all kingdoms of life. These oscillations control daily rhythms in metabolism, physiology and behavior, and they allow organisms to adapt their physiological needs to the time of day in an anticipatory fashion [1,2]. When the internal clocks run in synchrony with the external world, they provide organisms with significant competitive advantages [3].

Although the molecular clockwork components have widely divergent origins and are not conserved across the main divisions of life, a common design principle has been applied to all organisms where circadian timing mechanisms have been investigated. This paradigm relies on a negative transcription-translation feedback loop (TTFL), where protein products of clock genes feed back periodically to regulate their own expression and drive rhythmic output pathways and physiology [2,4,5]. Mounting evidence suggests that transcription-based oscillators are not the only means by which cells track time. Some examples of non-transcriptional oscillators are (i) the cyanobacterial phosphorylation oscillator, that can be reconstituted *in vitro* [6]; (ii) the circadian photosynthetic rhythms that persist in green algae after enucleation and hence in the absence of nuclear transcription [7]; or (iii) the peroxiredoxin (Prx) oxidation rhythms found in red blood cells [8], which do not have a nucleus.

Peroxiredoxins are a conserved family of antioxidant enzymes that maintain the cellular redox state by clearing the cell from reactive oxygen species (ROS). They reduce hydrogen peroxide (H_2_ O_2_) to water with the use of reducing equivalents provided by other physiological thiols [9]. As a result ROS removal, Prxs become oxidized in their active site [9]. Interestingly, and in contrast to the divergent evolution of the TTFL, levels of oxidized Prx have been shown to oscillate with a circadian period in all kingdoms of life [10]. The exact mechanism that generates circadian Prx redox rhythms, however, still remains an open question, and it seems to be different in different cell types (Figure 1A). Sulfiredoxin (Srx) is likely a key determinant of Prx3 hyperoxidation in adrenal gland, heart and brown adipose tissue [11] but not in red blood cells [12].

**Figure 1.**
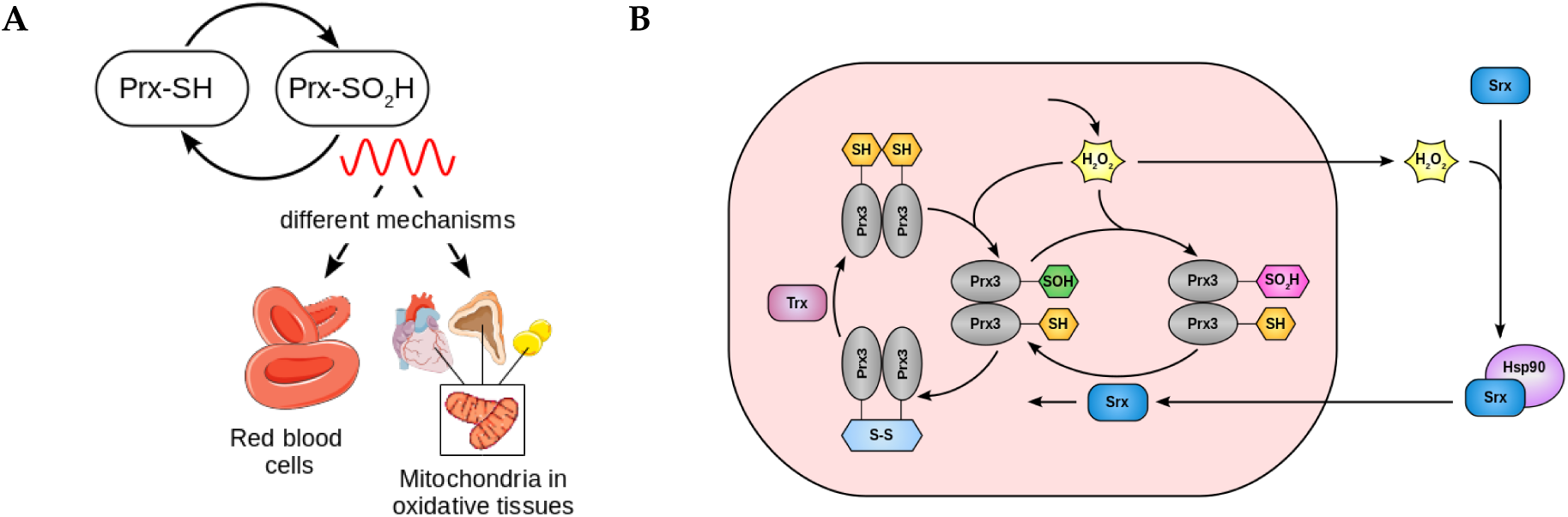
Peroxiredoxin oxidation cycles occur with circadian periodicity in all kingdoms of life. (**A**) The mechanisms for 24 h rhythm generation in Prx hyperoxidation in mammals differ in different cell types. **(B)** Model for the mechanism underlying circadian oscillation of Prx3-SO2H and Srx levels in mitochondria of adrenal gland, heart and brown adipose tissue. See text for details.

In this study, we use mathematical modeling to investigate the principles for Prx/Srx oscillations. Our results show that the combination of a fast Prx hyperoxidation event followed by a slow and delayed negative feedback loop is the minimal backbone that is required for the system to oscillate. We also find that this minimal motif produces relaxation-like oscillations with two temporal switch-like events, highlighting the importance of switches in the generation of oscillations.

## 2. Results

### 2.1 Derivation of a kinetic model for circadian redox oscillations

Because of their potential to oxidize and damage cellular protein and lipids, ROS levels must be under tight regulation. Prx3 is one of the major antioxidant proteins involved in H_2_O_2_ removal in mitochondria. A conserved cysteine (Cys) residue in its active site is oxidized by H_2_O_2_ to Cys-sulfenic acid (Prx3-SOH). This intermediate can react with a second conserved Cys from another Prx3 subunit to produce an intermolecular disulfide bond (Figure 1B), which can be reduced by the thioredoxin (Trx) system of the cell. Alternatively, Prx3-SOH can undergo further oxidation, termed hyperoxidation, in an S-sulfinylation reaction, to form a sulfinic acid (Prx3-SO_2_ H) [9,13]. Prx3-SO_2_ H is catalytically inactive [9,13]. It can be reduced back to Prx3-SOH by action of sulfiredoxin (Srx), which occurs for example in adrenal gland cells, brown adipocytes or cardiomyocytes (Figure 1B) [11]. In other cell types such as red blood cells, Prx-SO_2_ H has been described to get degraded [12].

In mitochondria from brown adipose, adrenal gland and heart tissue, Prx3-SO_2_ H and Srx levels oscillate with a circadian period [11]. H_2_O_2_ increase results in Prx3 inactivation, H_2_O_2_ accumulation and overflow to the cytosol. Cytosolic H_2_O_2_ activates pathways to control its own production. Among others, it stimulates Srx oxidation and intermolecular disulfide bridge formation with Hsp90 to promote translocation of Srx to the mitochondria, where it can reduce and thus reactivate Prx3-SO_2_ H [11]. Srx levels peak in mitochondria ∼8 h after the Prx3-SO_2_ H peak, and Srx becomes sensitive to degradation when Prx3-SO_2_ H levels decrease [11]. The mitochondrial import of Srx thus constitutes a negative feedback that enables a new cycle of H_2_O_2_ removal and Prx3-SO_2_ H accumulation (Figure 1B).

In order to understand how fast biochemical redox reactions result in slow 24 h rhythms, we develop a deterministic model containing the biochemical species from the Prx3/Srx system shown in Figure 1B. The complete set of equations of this large model, which we refer to as *detailed model*, are found in Appendix A. Unfortunately, quantitative details of the kinetic processes are not known and thus estimating parameters in such a large model represents a challenge. For this reason, and in order to gain insight into the design principles of redox rhythms in the Prx3/Srx system, we simplify the detailed model to its core motif (Figure 2A). Details of the model reduction are found in Appendix B.

**Figure 2.**
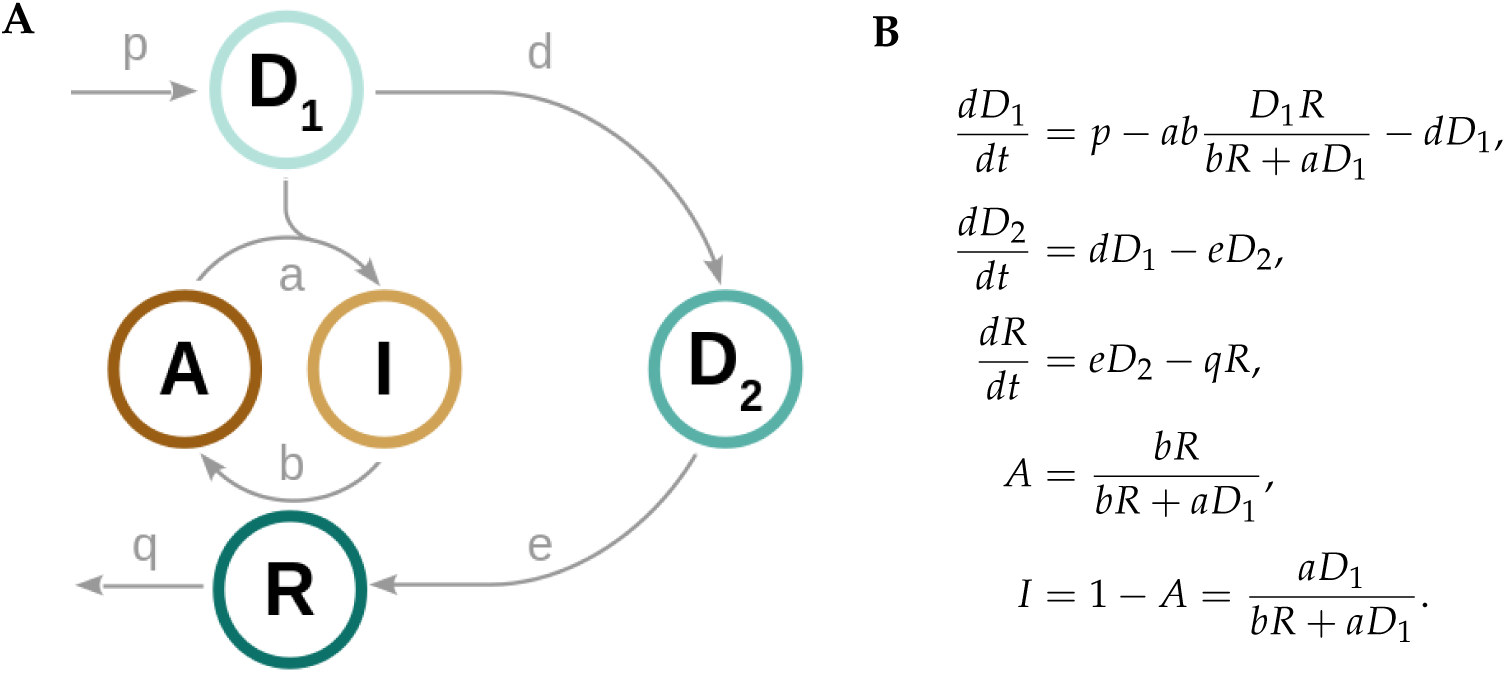
Core model for redox circadian oscillations. Scheme **(A)** and equations **(B)** of the core redox oscillator model. *A* represents active peroxiredoxin 3 (Prx3-SOH); *I*, inactive Prx3 (Prx3-SO_2_ H); *D*_1_, mitochondrial H_2_O_2_; *D*_2_, cytosolic H_2_O_2_ and *R*, mitochondrial Srx.

### 2.2 A low dimensional model represents the core Prx3-SO_2_ H/Srx circadian oscillator

We identify the core motif that is required for generation of Prx3-SO_2_ H and mitochondrial Srx circadian oscillations by systematically changing parameters and clamping variables to their mean value (Appendix B). The scheme and equations of this minimal model are shown in Figure 2. It only contains the variables, kinetic parameters and reactions that are necessary and sufficient for self-sustained rhythm generation.

The core oscillator model consists of only five variables and just one oxidation event (Figure 2A). *A* represents the *active* but partially oxidized Prx3 (Prx3-SOH) that gets further hyperoxidized by the *danger 1* (*D*_1_, mitochondrial H_2_O_2_) to produce the *inactive* Prx3-SO_2_ H, *I*. As the *A* →*I* reaction occurs, *A* levels decrease. The result is that the ability of mitochondria to remove *D*_1_ also decreases. When mitochondria can no longer eliminate the oxidant, *D*_1_ accumulates and overflows to the cytosol, where it is referred to as *danger 2* (*D*_2_, cytosolic H_2_O_2_). *D*_2_ activates the translocation of the *rescuer R* (mitochondrial Srx), that reduces *I* back to *A*, closing the negative feedback loop. The Prx3-SH oxidation to Prx3-SOH and the *D*_2_-induced oxidation of cytosolic Srx (Figure 1B) were found to be dispensable for rhythm generation. Thus these biochemical species as well as the reactions they take part in are omitted from the minimal model (details of the model simplification are provided in Appendix B).

We formulate ordinary differential equations (ODEs) to describe the behavior of the system over time. Production and removal terms are modeled with mass action kinetics (Figure 2B). We assume (i) constant *D*_1_ production (parameter *p* in the model) and (ii) constant Prx3 pool *A* + *I* over time; (iii) we apply the quasi-steady state approximation on *A* (Appendix B); and (iv) we only consider the negative feedback performed by *R*-induced reactivation of *A* stimulated by the *D*_2_ increase, although other additional feedback loops might exist, as seen by studies that have shown that cytosolic H_2_O_2_ *D*_2_ can activate pathways that directly decrease *D*_1_ production [14].

### 2.3 Design principle of the redox oscillator: fast A inactivation followed by a slow negative feedback loop

The dynamics of a system depend greatly on parameter values. Unfortunately, most kinetic rates in the Prx3/Srx sytem are not known in quantitative detail and therefore finding reasonable parameters represents a major challenge. Nevertheless, some physiological constraints can be taken into account to narrow down the plausible range of parameter values. First of all, the system needs to oscillate with a circadian period and the phase relationship between *I* and *R* should be approximately 8 h, according to previously published data [11]. Second, *in vitro* measurements of Prx3 hyperoxidation have determined a second order Prx3 hyperoxidation rate in the order of 10^4^ M^-1^ s^-1^ (*a* in the model), 1 000 – 10 000-fold higher than rate constants measured for cysteine thiols in other proteins [15]. A last constraint is derived from studies that have focused on H_2_O_2_ signaling and transport. H_2_O_2_ transport through biological membranes is thought to occur via aquaporins (AQP) [16]. The typical diameter of an aquaporin is ∼20 Å [17] and a study from 2011 showed that biological membranes typically contain 30 AQP/µm^2^ [18], although this number might vary across cell and membrane types. This implies that the total AQP area that can transport H_2_O_2_ out of the mitochondria is approximately 100 nm^2^ per µm^2^ of membrane. In other words, 0.01% of the membrane area contains AQP. Very generally, assuming that the probability of an H_2_O_2_ molecule to diffuse is in the same order of magnitude as its probability to get reduced by Prxs [19], but with the limitation that only 0.01% of the membrane allows H_2_O_2_ translocation, the probability of H_2_O_2_ being transported out of the mitochondria is 0.01% of its probability of getting reduced, i.e. 10 000-fold lower than its reduction rate *a*.

In order to obtain a reasonable set of parameters that satisfies the physiological constraints, we systematically vary all model parameters and perform bifurcation and sensitivity analyses. Bifurcation diagrams show under which parameter values the system oscillates and sensitivity analyses address how the oscillation period varies over the oscillatory range as a parameter is changed.

Bifurcation diagrams are presented for a selected choice of parameters in Figures 3A-C. The bifurcation plots show that the system oscillates for values of *d, e, q* < 1 (Figure 3A), where the limit cycle emerges at *d* = 0.02 and disappears at *d* = 0.5 (bifurcation plots for parameters *e* and *q* are shown in Appendix C). We also find that, in contrast to small *d, e, q* parameter values, the system requires a 1 000 – 10 000-fold higher *a* oxidation rate to enter into the oscillatory regime. The Hopf bifurcation occurs at *a* = 120 (Figure 3B) and once this value is reached, the oscillations persist for a wide range of kinetic parameters, with little effects in oscillation amplitude or period, indicating robustness of the model. The period sensitivity analyses reveal that the oscillation period depends strongly on the translocation rates *d* and *e* and on the *R* degradation parameter *q*, being most sensitive to changes in parameter *d* (Figure 3C and Appendix D).

**Figure 3.**
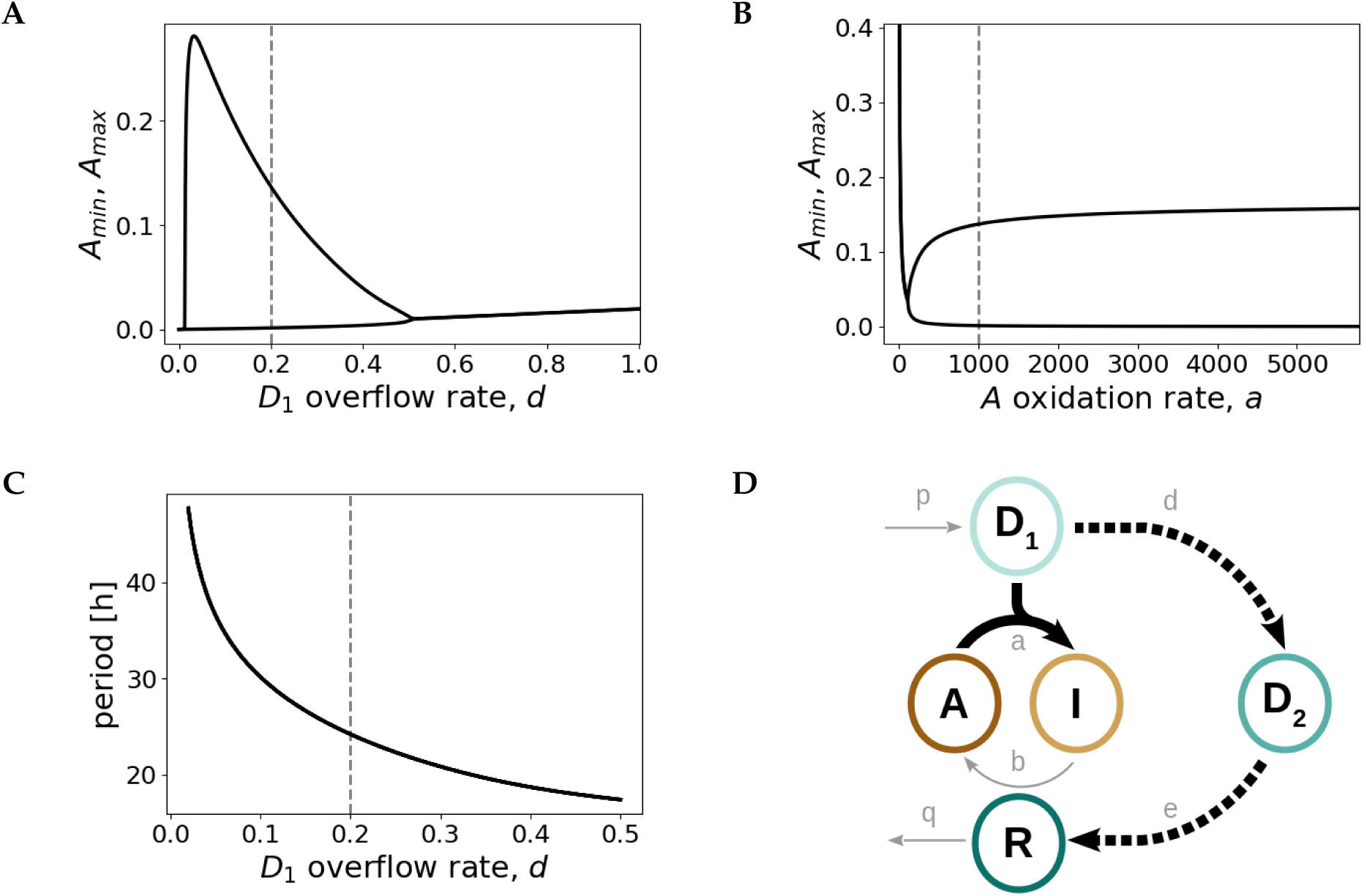
Fast *A* inactivation followed by a slow negative feedback loop is the design principle of the Prx3/Srx redox oscillator. Bifurcation diagrams as a function of the model parameters *d* **(A)** and *a* **(B)**. At *d* = 0.02 and *a* = 120 self-sustained oscillations emerge. *A*_*min*_ and *A*_*max*_ represent the minimum and maximum values of *A* in the oscillatory regime. **(C)** Variation of period as a function of *d*. The curves shown in panels **(A)**-**(C)** are obtained for the default parameter set given in the caption of Figure 4. The dashed gray lines depict the default parameter values given in the caption of Figure 4. **(D)** Sketch of the core backbone for *I*/*R* redox oscillations: fast *D*_1_-induced *A* inactivation (continuous thick line) followed by a slow *D*_1_ *- D*_2_ *- R* negative feedback loop (dashed line).

These findings show that a fast *A* inactivation reaction (high *a* value) in combination with a slower negative feedback loop (1 000 – 10 000-fold lower *d, e, q* rates) constitute the backbone of the redox oscillator model (Figure 3D). We thus find a plausible set of model parameter values that produce robust circadian oscillations with the expected characteristics: a period of 24.2 h and a *I*/*R* phase difference of 8.7 h, as determined by maxima estimation of the *R* and *I* dynamics (Figure 4). Moreover, the parameter choice is in agreement with the high Prx3 oxidation rate measured experimentally [15] and with the ∼10 000-fold slower physiological H_2_O_2_ translocation.

**Figure 4.**
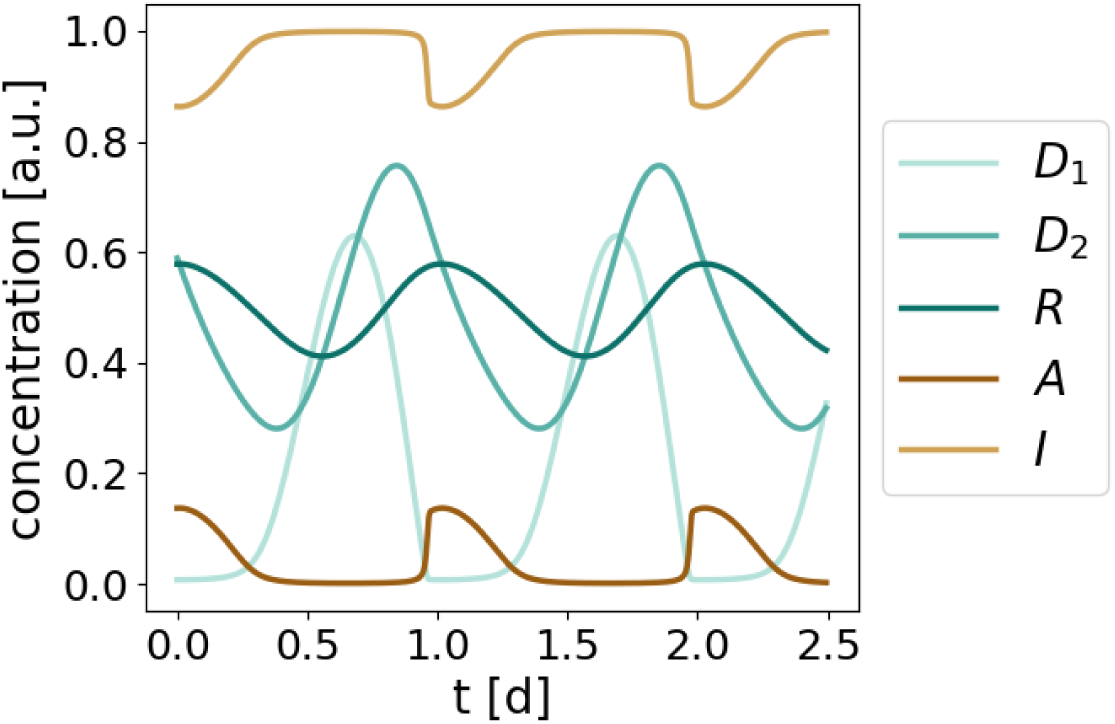
Dynamics of the redox model. Limit cycle oscillations obtained by numerical integration of the equations shown in Figure 2B for the following parameter values (arbitrary units, a.u.): *p* = 1, *a* = 1 000, *b* = 2, *d* = 0.2, *e* = *q* = 0.1. Consistent with experimental data [11] and with the model constraints, the resulting period is 24.2 h and the phase shift between *I* and *R* is 8.7 h.

According to our results, only a small fraction of active Prx3 *A* is needed to keep *D*_1_ levels close to 0, as seen by the higher *I* levels compared to *A* during the whole simulated time (Figure 4). The results also predict that *D*_2_ levels peak 3.9 h after *D*_1_ (or 4.5 h after *I*), an observation that could be relevant in the context of the canonical TTFL and will be further addressed in the Discussion section.

Interestingly, insights from the redox oscillator model apply as well to larger and more complex models. Firstly, negative feedback loops (*D*_1_ *- D*_2_ *- R* in the minimal model) as well as nonlinear terms (given by the *A* equation, Figure 2B) are required to achieve self-sustained oscillations [20]. Secondly, overcritical delays about quarter to half of a period are necessary for oscillations to arise [20–22] (in the minimal model, slow *d, e, q* rates are required to yield the 8.7 h delay). And lastly, translocation and degradation rates have profound effects on the period [23–26]. It should be noted, nevertheless, that the choice of parameters is not tailored to specific kinetic data. Kinetic parameters might differ among tissues and might also depend on physiological conditions.

### 2.4 The I/R redox oscillator is characterized by three phases and two temporal switches

We have shown that, in order to oscillate, the mitochondrial Prx3-SO_2_ H/Srx (*I*/*R*) system requires a fast *D*_1_-induced inactivation of *A* followed by a lengthy and slower negative feedback loop *D*_1_ *- D*_2_ *- R* that reactivates *A* (Figure 3D). As seen by the equations from Figure 2B, the dynamics of *A, I* and *D*_1_ depend on the fast *a* oxidation rate compared to the slower dynamics of *D*_2_ or *R*. This is reflected in the relaxation-or triangular-like waveforms of *A, I* and *D*_1_ (Figure 4). Such relaxation oscillations are characterized by two consecutive processes that occur on different timescales. There are intervals of time, during which little happens (*A* = 0 or *I* = 1 in the system’s dynamics), followed by time intervals with considerable changes. These oscillators produce the triangular-like dynamics observed for *A, I* and *D*_1_ (Figure 4).

It is known that oscillations depend on negative feedbacks, but that in the absence of appropriate delays or nonlinear terms, negative feedback circuits often settle into a stable steady state termed homeostasis [20,27,28]. Several computational studies have demonstrated that sufficiently strong nonlinearities are required to generate self-sustained oscillations [26,29–31]. Such nonlinear terms often exhibit sigmoidal or *switch-like* characteristics. Surprisingly, no switch-like curve is evident from the only nonlinear term of the model 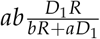 (Figure 2B and Appendix B), nor from the response of any variable to upstream components (data not shown). Nonetheless, we find that this term is at the core of the oscillations, as its linearization results in the system settling into a stable steady state. The large value of *a* allows the production of self-sustained rhythms, even in the absence of an explicit kinetic switch.

In order to emphasize the importance of the nonlinear term in the context of oscillations, and given the switch-like dynamics of *A, I* and *R*, we still use the term *switch*. Instead of using it to describe how the steady state of a first component responds to a second component which is upstream of the former (often done in the literature [31–34]), we focus here on temporal switches that mark transitions between phases. We see that the model dynamics can be split into three phases separated by two temporal switches (Figure 5).

**Figure 5.**
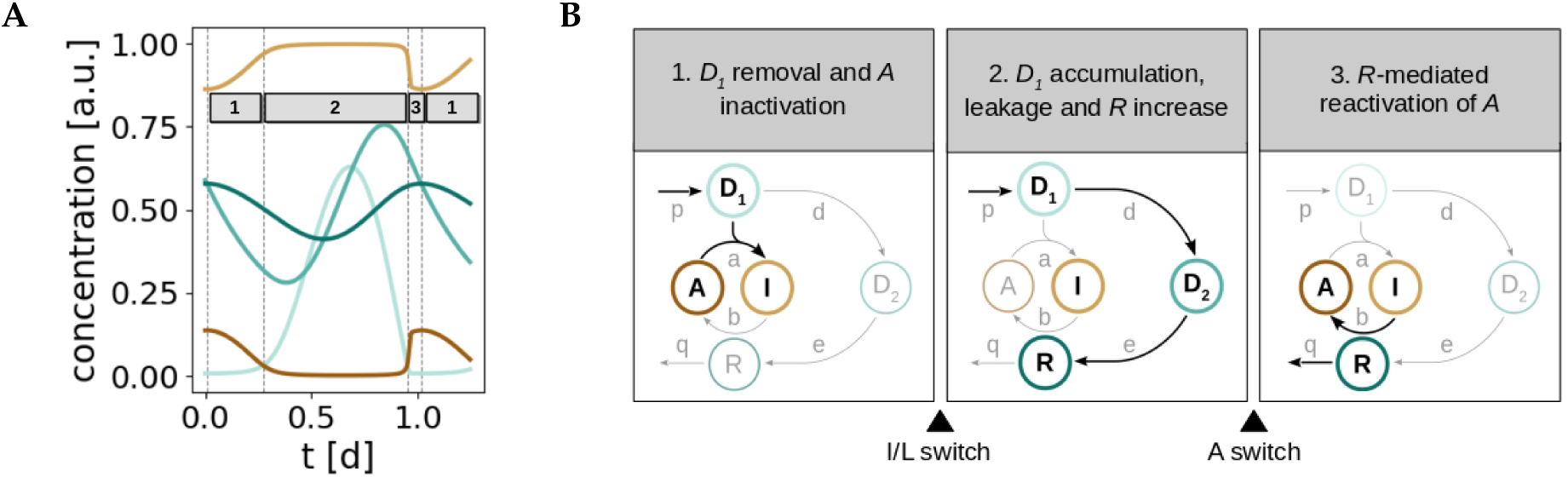
A series of three phases and two temporal switches can be regarded as the mechanism for generation of redox oscillations. **(A)** Division of the temporal dynamics of the *I*/*R* system into three phases that explain the major transitions of the model components. **(B)** Scheme of the major events that take place in each phase (see main text for details).

The first phase is characterized by the inactivation of *A* as a result of *D*_1_ removal. As long as there is active Prx3 (*A*) in the system, mitochondrial H_2_O_2_ (*D*_1_) levels are kept in check and *D*_1_ is removed. But the cost of the *D*_1_ clearance is the inactivation of *A* to *I*, and thus *A* levels decrease as *I* increases. When most of the active Prx3 pool *A* has been hyperoxidized and inactivated to *I*, the first switch occurs and the system progresses to the next phase. As a consequence of the *A inactivation*, its ability to remove *D*_1_ molecules decreases and thus *D*_1_ accumulates and *leaks* to the cytosol. In other words, the flux of *D*_1_ changes from inactivating *A* to leaking, what gives this first temporal switch its name: *inactivation/leakage* (I/L) switch. As a result of the I/L switch, *D*_1_ accumulates, leaks and activates the *D*_1_ *- D*_2_ *- R* negative feedback. Consequently, *D*_2_ and *R* levels increase (phase 2). At the end of the second phase, the increase in *R* is accompanied by a decrease in *D*_1_. Once *D*_1_ and *R* reach critical values in the nonlinear term, the negative feedback becomes effective and the dynamics of *A*, that are governed by the nonlinearity (Figure 2B), change suddenly. This triggers the *activation* (A) switch and the sharp increase in *A* levels (phase 3). The system is thus switched to its active state and a new round of *D*_1_ removal can start. Furthermore, *R* becomes sensitive to degradation. Our series of two temporal switches separate the two timescales in the relaxation oscillations: the I/L switch sets the start of the slow timescale (the negative feedback loop), whereas the A switch marks the beginning of the fast clearance of *D*_1_.

## 3. Discussion

We have designed the first model for the complex biochemical system of Prx3-SO_2_ H/Srx redox oscillations and we find that the loop Prx3-SOH – Prx3-SO_2_ H (Figure 1B) is necessary and sufficient for the generation of oscillations. The design principles of this oscillator are (i) a fast Prx3-SOH inactivation followed by (ii) a slow and delayed negative feedback loop where mitochondrial H_2_O_2_ leaks to the cytosol to promote the translocation of cytosolic Srx to mitochondria, approximately 9 h after the inactivation of Prx3-SOH. This simple backbone reproduces the previously described circadian oscillations of mitochondrial Prx3-SO_2_ H and Srx as well as the delay in mitochondrial Srx import.

It is known that relaxation oscillations typically depend on positive feedback loops [35,36]. However, even in the absence of explicit positive feedback loops, the redox oscillator model still produces oscillations of Prx3-SOH, Prx3-SO_2_ H and mitochondrial H_2_O_2_ that are of relaxation type. Two temporal switches characterize their triangular-like waveform and divide the system’s dynamics into three phases. A summary is shown in Figure 6. This scheme with three phases and two switches is reminiscent to that of the cell cycle. A series of biochemical switches control transitions between the various phases of the cell cycle. They maintain its orderly progression and act as checkpoints to ensure that each phase has been properly completed before progression to the next phase. Such switches have been shown to generate decisive, robust (and potentially irreversible) transitions and trigger stable oscillations [28,37,38]. It should be noted, however, that the switches that regulate cell cycle transitions are the consequence of a complicated network of positive and negative feedback loops and result in bistability, as opposed to our simple model that (strikingly) contains only one negative feedback loop. The redox model highlights the importance of negative feedbacks and the diversity of switches in the context of oscillations.

**Figure 6.**
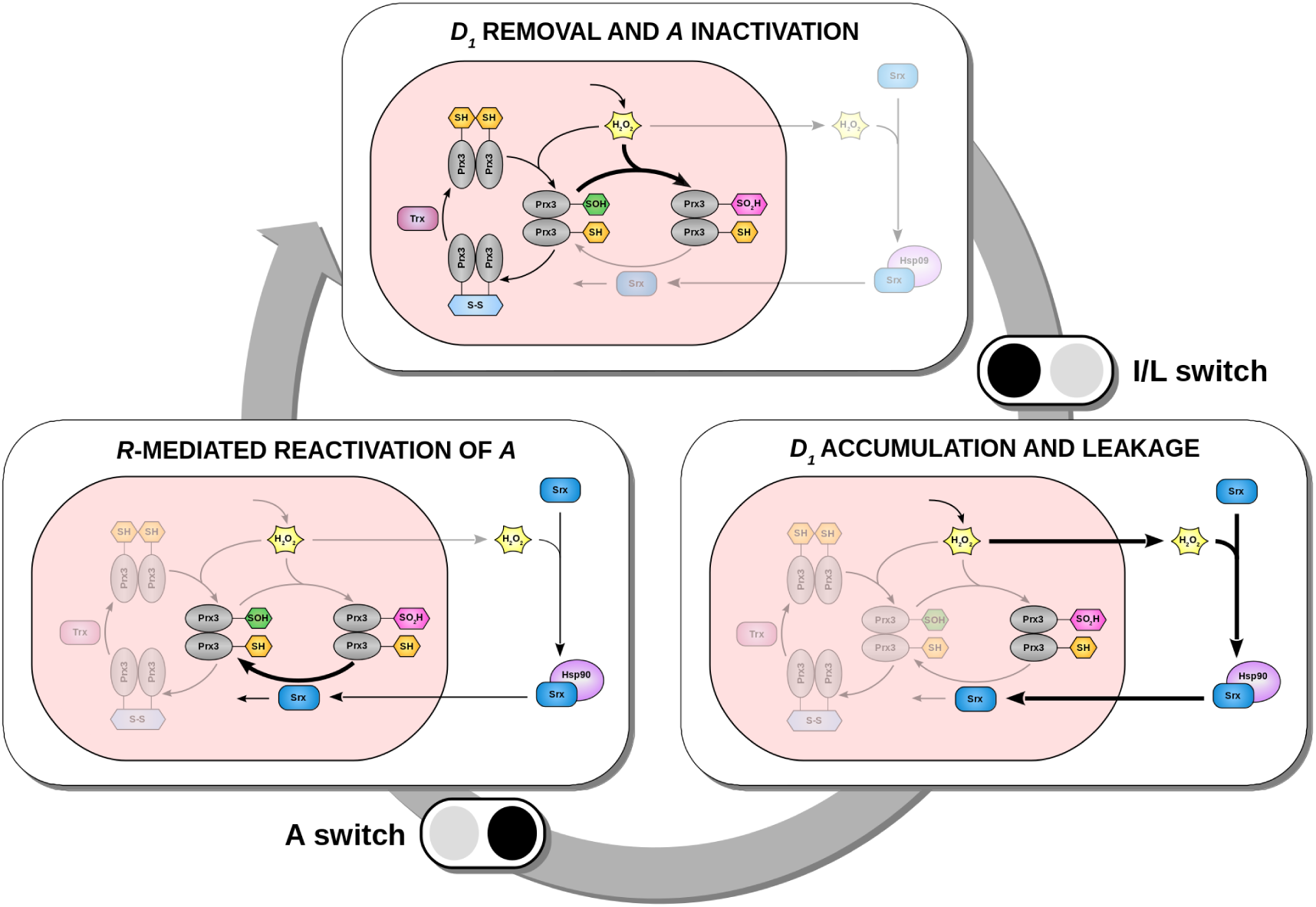
Model for the mechanism underlying circadian oscillations of Prx3-SO_2_ H and Srx in mitochondria of mammalian adrenal gland, heart and brown adipose tissue. Three phases and two temporal switches conceptually explain the generation of oscillations. The switches are represented by the black/gray icons. The switch occurs from black to gray: the black circle represents the phase from which the system comes, and the gray circle depicts the phase it progresses to.

Although quantitative details of the kinetic parameters in the Prx3/Srx system remain largely unknown, we have found a reasonable set of parameters that is consistent with the biology they describe. Previous biochemical studies have estimated that Prxs are 1 000 – 10 000 times more reactive to H_2_O_2_ than other reduced cellular thiols [15], and that thus they can confine the diffusion of the oxidant [19]. Under the assumption that 0.01% of the membrane area contains transport proteins (AQP) that can passively transport H_2_O_2_ out of the compartment [18], the translocation rate of H_2_O_2_ is 10 000-fold smaller than its reduction rate. This is in agreement with the parameter values, *a* = 1000 and *d* = 0.2. Since the import of Srx to mitochondria also requires a protein transporter, we have assumed this rate to be in the same order of magnitude as the translocation of H_2_O_2_ to the cytosol. The experimental determination of the rate at which a protein is transported through a membrane is challenging, because the transport depends on a number of factors including pH, membrane voltage, protein length, protein 3D conformation, membrane fatty acid composition, and ATP levels (if the transport is active), among others. Hence the estimation of translocation kinetics is only preliminary. Furthermore, the choices for Srx degradation rate *q* and hyperoxidized Prx3 reduction rate by Srx *b* are consistent with previous experimental studies, which have estimated a half-life of sulfiredoxin of 4-5 h [11,39], and a rate of reduction of hyperoxidized Prx by Srx of approximately 2 M^-1^ s^-1^, 1 000 – 10 000 times smaller than its oxidation rate [40].

Also the detailed model (Appendix A) oscillates along the lines of the design principles identified for the reduced model. It shows the same characteristics: relaxation oscillations, temporal switches and expected period and *I*/*R* phase delay, supporting the robustness of the minimal model. According to biochemical assays, the rate constant for Prx3-SH oxidation to sulfenic acid Prx3-SOH is in the order of 10^7^ M^-1^ s^-1^ [15]. From the simulations of the detailed model we see that when this rate (*k*_1_ in the model) increases, the amplitude of Prx3-SO_2_ H and Srx oscillations also increases (Appendix A). This suggests that although the loop Prx3-SH – Prx3-SOH – Prx3-SS – Prx3-SH is not required for oscillations, it fine-tunes the dynamics of the Prx3/Srx oscillatory system. A high Prx3-SO_2_ H and Srx amplitude is in agreement with experimental data. Western blots show intense bands at the peak of Prx3-SO_2_ H or Srx oscillations and no bands at the trough [11].

The modeling approach described in this study assumes constant H_2_O_2_ production and no feedback regulation from cytosolic H_2_O_2_ to reduce mitochondrial production. However, recent experimental work has shown that (i) steroidogenesis in adrenal gland (a major source of H_2_O_2_) is under the control of the circadian TTFL [41,42] and that (ii) cytosolic H_2_O_2_ can signal as a second messenger and activate the p38 MAPK pathway to decrease H_2_O_2_ production [11,14]. It is also becoming evident that the cellular redox poise can feed back to the TTFL oscillator [43–45]. The H_2_O_2_ leakage might represent a potential coupling node between both clocks. Our model predicts that H_2_O_2_ peaks in the cytoplasm ∼4.5 h after the peak of Prx3-SO_2_ H (or ∼4 h after mitochondrial H_2_O_2_). This observation might represent an attractive target for redox sensors that could measure when H_2_O_2_ peaks in the cytosol in relation to its peak in mitochondria. The peak of cytoplasmatic H_2_O_2_ at a proper time might be of importance to transmit appropriately timed signals about the redox state of the cell to the canonical TTFL oscillator. It is likely that a stable phase relationship is required between both oscillators for their optimal function.

It should be noted that the redox model reproduces the circadian rhythms at the level of a *single* mitochondrion. However, depending on the cell type, mitochondria vary in number. For example, a human liver cell contains 1 000 – 2 000 mitochondria, whereas a cardiac myocyte contains 7 000 – 10 000 mitochondria [46]. Although single mitochondria might be competent redox oscillators, they could still synchronize their circadian cycles to each other. It seems plausible that coupling between mitochondria is needed in order to transmit the information about the redox state of the cell to the TTFL clock as an ensemble.

Finally, we close with a short remark about the beauty of simple and generic models. We have elaborated on the basic ingredients that constitute the Prx3/Srx redox oscillator in mammalian mitochondria from adrenal gland, heart and brown adipose tissue. We have found the core motif that results necessary and sufficient to generate self-sustained rhythms that reproduce previously published experimental data. Our model has been implemented using plausible and simple, yet generic mathematical representations. This five-variable network might be relevant for other simplified oscillatory systems that follow the same logic of fast (in)activation followed by slow negative feedback, such as the mitotic oscillator involving cyclin and Cdc2 kinase interactions or the Ca^2+^ oscillations based on Ca^2+^-induced Ca^2+^ release [47]. The fact that nature has converged on common mathematical structures underlines the links between similar dynamic phenomena occurring in widely different biological settings. It even constitutes a step forward in the artificial design of reliable biological clocks [48,49].

## 4. Materials and Methods

All temporal simulations and model analyses have been performed with Python, using the odeint integrator from the scipy module.

## Author Contributions

Conceptualization, Marta del Olmo and Hanspeter Herzel; Data curation, Marta del Olmo and Hanspeter Herzel; Formal analysis, Marta del Olmo and Hanspeter Herzel; Funding acquisition, Achim Kramer and Hanspeter Herzel; Investigation, Marta del Olmo and Hanspeter Herzel; Methodology, Marta del Olmo and Hanspeter Herzel; Project administration, Achim Kramer and Hanspeter Herzel; Resources, Marta del Olmo and Hanspeter Herzel; Software, Marta del Olmo and Hanspeter Herzel; Supervision, Achim Kramer and Hanspeter Herzel; Validation, Marta del Olmo and Hanspeter Herzel; Visualization, Marta del Olmo and Hanspeter Herzel; Writing – original draft, Marta del Olmo and Hanspeter Herzel; Writing – review & editing, Achim Kramer and Hanspeter Herzel.

## Acknowledgments

We are grateful for fruitful discussions with Dr. Bharath Ananthasubramaniam and Abhishek Upadhyay.

## Abbreviations

The following abbreviations are used in this manuscript:

*A*: Active Prx3 (Prx3-SOH)
A: Activation
AQP: Aquaporin
a.u.: Arbitrary unit
Cys: Cysteine
d: Day
*D*_1_: Danger 1 (mitochondrial H_2_O_2_)
*D*_2_: Danger 2 (cytosolic H_2_O_2_)
h: Hour
H_2_: O Water
H_2_O_2_: Hydrogen peroxide
*I*: Inactive Prx3 (Prx3-SO_2_ H)
I/L: Inactivation-Leakage
MAPK: Mitogen-activated protein kinase
ODE: Ordinary differential equation
Prx: Peroxiredoxin
*R*: Rescuer (Srx)
ROS: Reactive oxygen species
Srx: Sulfiredoxin
Trx: Thioredoxin
TTFL: Transcription-translation feedback loop

## Appendix A. Detailed model of redox oscillations

We develop a deterministic model that contains all biochemical species from the Prx3/Srx oscillating system shown in Figure 1B [11]. A scheme of the model and its equations are shown in Figure A1A-B. Production and degradation terms are modeled assuming mass action kinetics. The total Prx3 pool (*SH* + *SOH* + *SO*_2_ *H* + *SS* in the model) and the production of H_2_O_2_ (*k*_0_ in the model) are assumed to be constant over time.

Applying the two ingredients that we identify as design principles of the oscillator, namely fast *SOH* inactivation followed by slow *H*_2_ *O*_2*M*_ *- H*_2_ *O*_2*C*_ *- Srx*_*OX*_ *- SrxHsp - Srx*_*M*_ feedback, we obtain oscillations in all 11 variables. The oscillations of the 5 core variables that comprise the minimal oscillator are shown in Figure A1C. The resulting period is 23.5 h and the phase difference between Prx3-SO_2_ H and mitochondrial Srx is 7.3 h. These simulations support the robustness of the minimal model, since applying its design principles into a larger model also generates oscillations with similar characteristics.

Parameters are chosen according to the constraints that we use to infer the parameter values from the minimal model. Degradation and translocation rates are kept in the order of magnitude of their corresponding parameters on the minimal model (*d, e, q*). We set *k*_7_ (the rate of oxidation of cytosolic Srx that results in an intermolecular bridge with Hsp) to 2, following previous results, which have estimated that Prxs react 1 000 – 10 000 times faster with H_2_O_2_ than other reduced cellular thiols [15]. Biochemical assays have determined the rate constants for Prx3-SH oxidation to sulfenic acid Prx3-SOH (*k*_1_ in the detailed model) and hyperoxidation to Prx3-SO_2_ H (*k*_2_ in the detailed model) to be in the order of 10^7^ M^-1^ s^-1^ and 10^4^ M^-1^ s^-1^, respectively [15]. We observe that the amplitude of Prx3-SO_2_ H and mitochondrial Srx oscillations increases as *k*_1_ is increased, a result that reproduces the details from published western blot data. Kil and colleagues have shown in [11] intense bands for both Prx3-SO_2_ H and mitochondrial Srx at their peak times and no bands at their troughs, implying that the oscillation amplitude is large.

**Figure A1.**
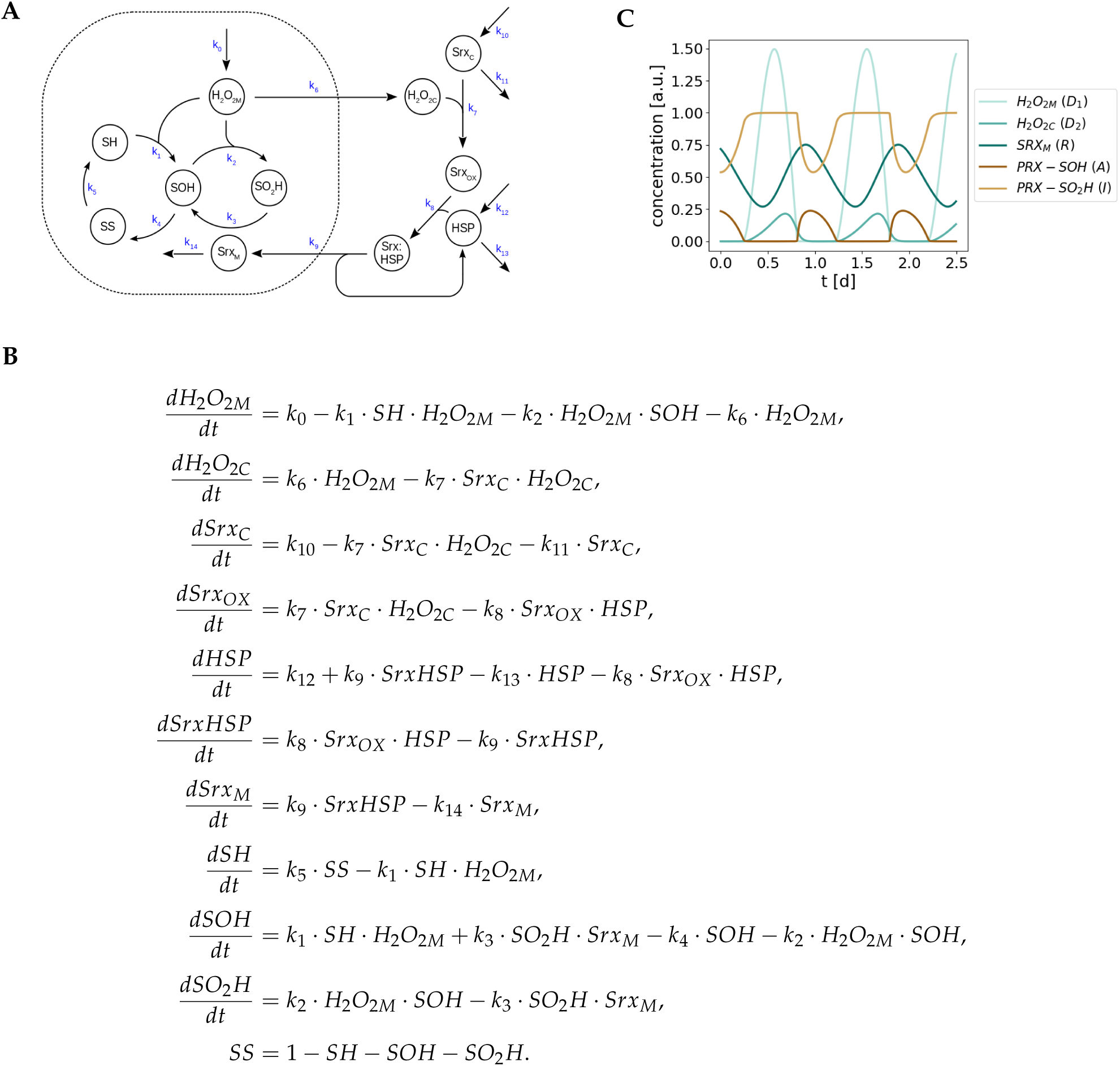
Detailed 10-ODE model for circadian redox oscillations. (**A**) Scheme of the biochemical species identified in the mitochondrial Prx3-SO_2_ H/Srx circadian oscillating system. **(B)** Equations of the model. **(C)** Limit cycle oscillations obtained by numerical integration of the equations shown in (B) for the following parameter values (a.u.): *k*_0_ = 1, *k*_1_ = 10^7^, *k*_2_ = 10^4^, *k*_3_ = 2, *k*_4_ = 1, *k*_5_ = 1, *k*_6_ = 0.2, *k*_7_ = 2, *k*_8_ = 0.5, *k*_9_ = 0.2, *k*_10_ = 0.2, *k*_11_ = 0.1, *k*_12_ = 0.2, *k*_13_ = 0.1, *k*_14_ = 0.2. The period of oscillation is 23.5 h; the phase shift between Prx3-SO_2_ H and mitochondrial Srx is 7.3 h.

## Appendix B. Model simplification

The minimal redox oscillator model used in this paper (Figure 2) is a condensed representation of the complex biochemistry of the Prx3/Srx oscillating system in mitochondria from adrenal gland, heart and brown adipose tissue (Figure 1B). Whereas the detailed model contains all 11 variables from Figure 1B (Appendix A), the minimal model is a relatively small network with only 5 variables. Because the detailed model contains 15 parameters, most of which are unknown, obtaining oscillations is challenging. For this reason, the 10-ODE detailed model is simplified to the 6-ODE model shown in Figure A2. Apart from the 5 variables from the minimal model, *S* represents cytosolic Srx, *A*_1_ represents the Prx3 in its reduced thiol form (Prx3-SH), and *A*_2_ represents the sulfenic acid Prx3-SOH. Again, mass action kinetics is used to model production and degradation terms and we assume constant mitochondrial H_2_O_2_ production *p* as well as constant Prx3 pool (*A*_1_ + *A*_2_ + *I*) over time.

We find oscillations for the parameter set shown in the caption of Figure A2. To gain insight into the design principles of the Prx3/Srx oscillator and in line with previous theoretical studies [50], we systematically clamp all seven variables. To clamp a variable or a process means to set it to its mean (constant) value, trying to resemble conditions of constitutive expression from the wet lab. This strategy, instead of simply removing a variable, allows to compare the effect of rhythmic versus basal regulation. Thus, if a state variable is fixed at its mean level (i.e., clamped) and the network remains rhythmic, this means that the clamped variable is not required for the generation of oscillations. It might play a role in regulating the oscillation period or amplitude, but it is not necessary to obtain self-sustained oscillations. By systematically clamping all 7 variables we find that neither *A*_1_ nor *S* are necessary for the generation of self-sustained oscillations and thus the 6-ODE model from Figure A2 is simplified to the model shown in Figure A3A. This model version contains the 5 core variables from the minimal model described by 4 ODEs, 6 parameters and 2 bilinear reactions (equations are shown in Figure A3B). It oscillates with a 25.3 h period and the phase difference between Prx3-SO_2_ H and mitochondrial Srx is 8.3 h (A3C), as expected.

Interestingly, and as shown in Figure A3D, we observe that the production of *A* (defined by the bilinear term *bIR*, Figure A3B) is very similar to its removal (defined by the second bilinear term *aAD*_1_) during the whole simulated time. Therefore, and in order to further simplify the model, we assume *aAD*_1_ = *bIR* over time (quasi-steady state assumption). This allows the replacement of the *A*-ODE by the following algebraic equation:

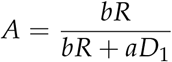

This is the equation in the minimal model described in the main text and shown in Figure 2B. The two bilinear terms, due to the quasi-steady state approximation, are thus coupled and transformed into the only nonlinear term in the minimal model (Figure 2B). The dynamics of the system do not change significantly (compare oscillations from Figures 4 and A3C).

The fact that we can introduce the quasi-steady state approximation on A tells much about the system. In general, the quasi steady state assumption is used when one part of the system reacts much more quickly than another [51,52]. The module that reacts faster is often said to have a shorter characteristic timescale or to equilibrate faster than the slower one. And in chemical terms, when something equilibrates fast, it can be said that it is in steady state with respect to the slower moving system. The time dependence of the faster reacting module can be simplified, reducing the number of variables that the system has to be solved for. In the context of the Prx3/Srx oscillating system, the quasi-steady state approximation performed on *A* means that *A* and *I* react faster than the rest of the variables, and this can indeed be seen in their sharp changes and in the two switches that occur over time (Figure 5B).

**Figure A2.**
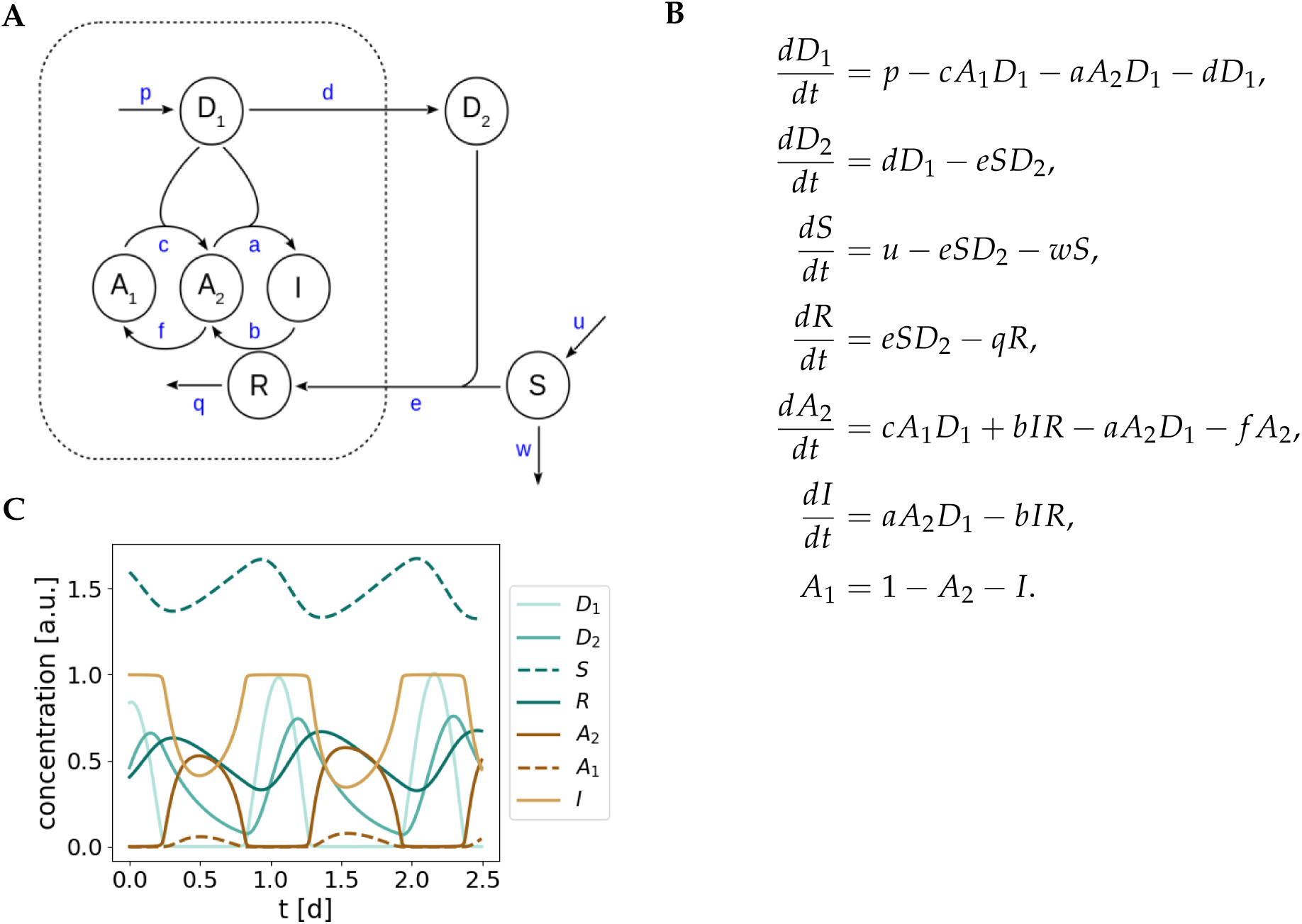
6-ODE model for circadian redox oscillations. (**A**) The 10-ODE detailed model is simplified to the 7-variable model shown in this figure. *A*_1_ represents Prx3-SH; *A*_2_, Prx3-SOH; *I*, Prx3-SO_2_ H; *D*_1_, mitochondrial H_2_O_2_; *D*_2_, cytosolic H_2_O_2_; *S*, cytosolic Srx and *R*, mitochondrial Srx. (**B**) Equations of the 7-variable model. We assume constant *A*_1_ + *A*_2_ + *I* over time and thus the number of ODEs is reduced to 6. The dynamics of *A*_1_ are described by a mass conservation law. (**C**) Limit cycle oscillations obtained by numerical integration of the equations shown in (B) for the following parameter values (a.u.): *a* = 1000, *b* = 2, *c* = 10000, *d* = *u* = 0.2, *e* = *q* = *w* = 0.1, *f* = *p* = 1. The period of oscillation is 25.3 h; the phase shift between *I* and *R* is 8.3 h.

**Figure A3.**
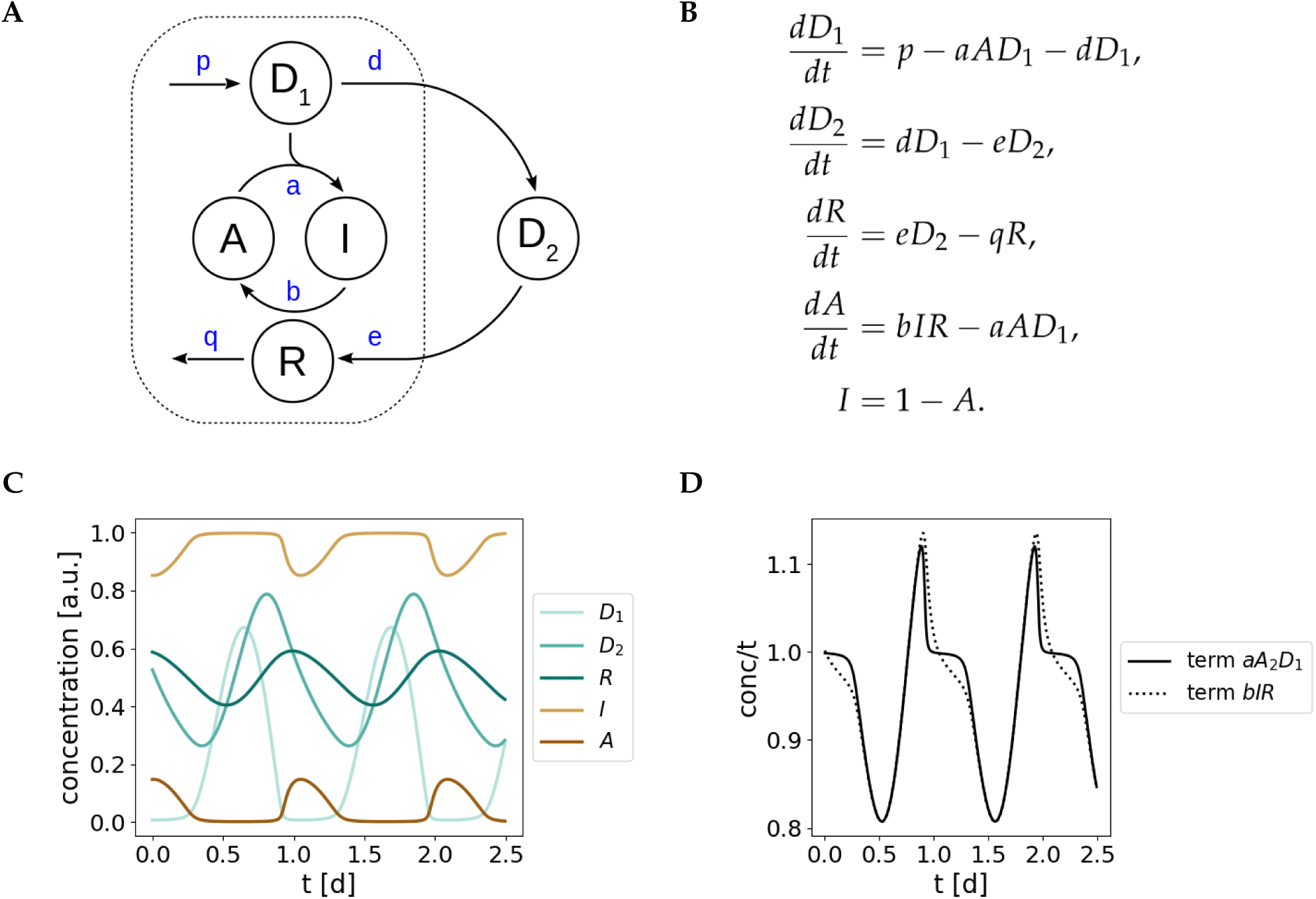
Reduced 4-ODE model for circadian redox oscillations. (**A**) The 6-ODE model is simplified to the 5-variable model shown in this figure. *A* represents Prx3-SOH; *I*, Prx3-SO_2_ H; *D*_1_, mitochondrial H_2_O_2_; *D*_2_, cytosolic H_2_O_2_ and *R*, mitochondrial Srx. (**B**) Equations of the 5-variable model. We assume constant *A* + *I* over time and thus the number of ODEs is reduced to 4, with the dynamics of *I* being described by a mass conservation law. (**C**) Limit cycle oscillations obtained by numerical integration of the equations shown in (B) for the following parameter values (a.u.): *a* = 1000, *b* = 2, *c* = 10000, *d* = 0.2, *e* = *q* = 0.1, *p* = 1. The period of oscillation is 24.5 h; the phase shift between *I* and *R* is 8.8 h. (**D**) Similar evolution of the bilinear terms *bIR* and *aAD*_1_ over time for the parameter values in (C), allowing the quasi-steady state state approximation that leads to the simplified model shown in Figure 2B.

## Appendix C. Bifurcation analyses

We vary all model parameters and perform bifurcation analyses to obtain a plausible set of parameters that satisfy the physiological constraints. Bifurcation diagrams for the Srx translocation rate (parameter *e*), mitochondrial Srx degradation rate (*q*), Prx3-SO_2_ H reduction rate by Srx (*b*) and mitochondrial H_2_O_2_ production (*p*) are shown in Figure A4. A note on units: the first order translocation and degradation rates *e* and *q* have units of h^-1^; the second order reduction rate *b* needs inverse units of time (h^-1^) and inverse units of concentration (arbitrary) and *D*_1_ production rate *p* has units of concentration (arbitrary) divided by time (h).

**Figure A4.**
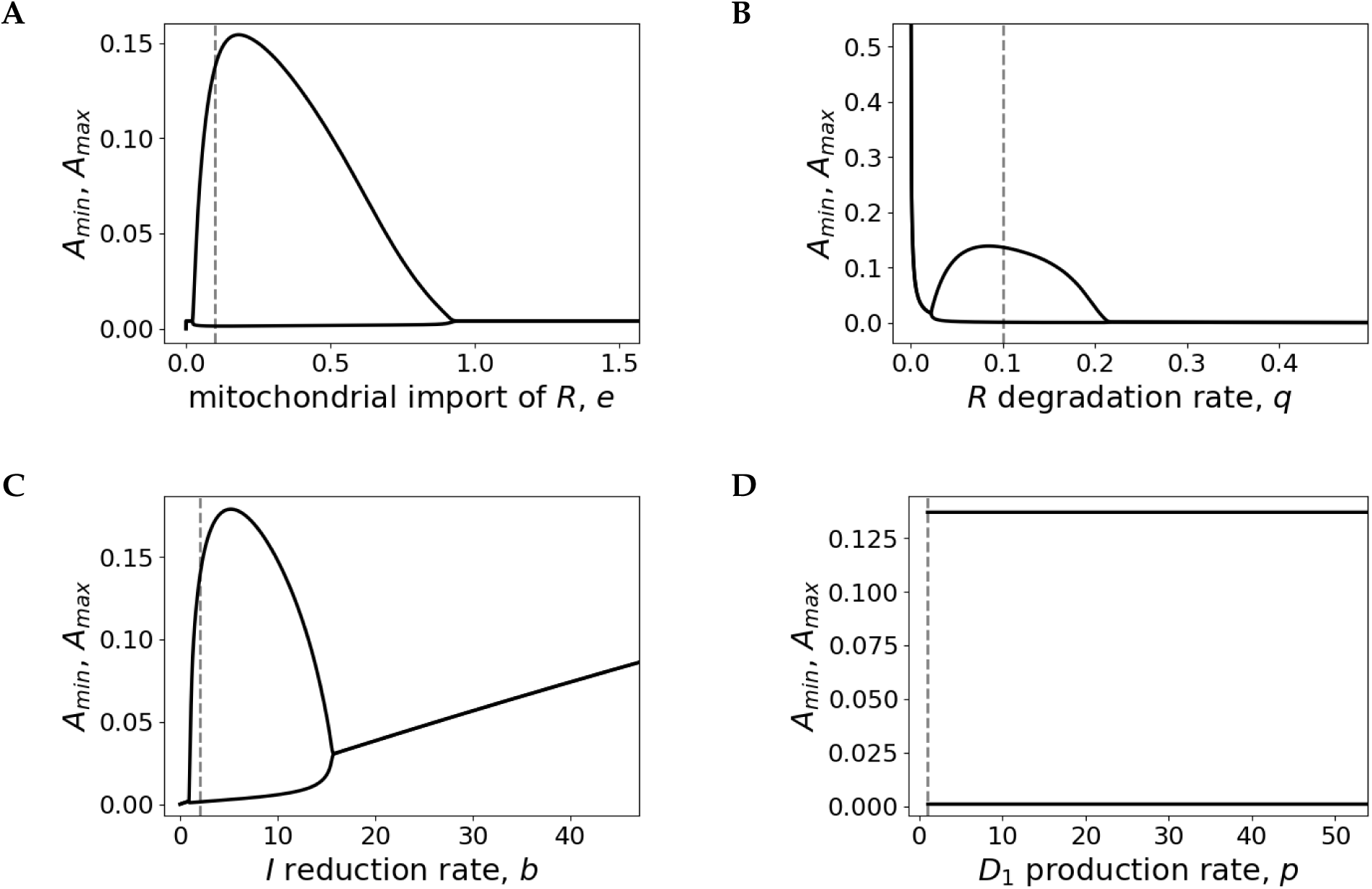
Bifurcation diagrams as a function of the model parameters *e* **(A)**, *q* **(B)**, *b* **(C)** and *p* **(D)**. At *e* = 0.03, *q* = 0.025 and *b* = 1 self-sustained oscillations emerge. The system oscillates for any positive value of *p*. *A*_*min*_ and *A*_*max*_ represent the minimum and maximum values of *A* in the oscillatory regime. The curves are obtained for the default parameter set shown in the caption of Figure 4, by numerical integration of the model equations given in Figure 2B. The dashed gray lines refer to the default parameter values.

## Appendix D. Period sensitivity analyses

We also analyze how sensitive the behavior of the redox oscillator model is towards parameter changes. We increase and decrease all model parameters by 10% and quantify the effect of such change on the oscillation period. The results are shown in Figure A5. As expected [25,26], translocation and degradation rates *d, e* and *q* control the period most strongly, but interestingly also the reactivation kinetics represented by *b* has some effects on the period.

**Figure A5.**
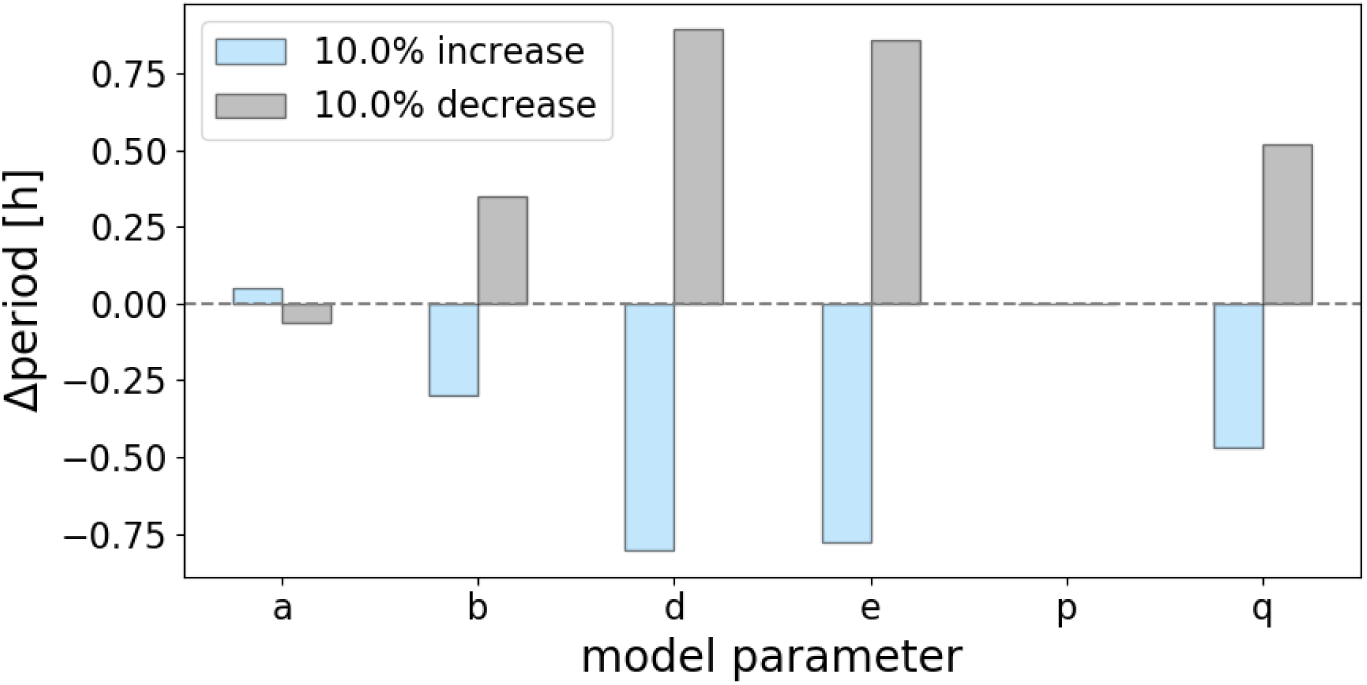
Period sensitivity analysis as a function of a 10% increase (blue) or 10% decrease (gray) of the default parameter value shown in the caption of Figure 4. Δperiod [h] represents the difference between the period calculated from the new parameter set and the period from the default parameter set. Period values are obtained for each parameter set by numerical integration of the model equations given in Figure 2B.

